# Probabilistic axon targeting dynamics lead to individualized brain wiring

**DOI:** 10.1101/2022.08.26.505432

**Authors:** Maheva Andriatsilavo, Alexandre Dumoulin, Suchetana Bias Dutta, Esther T. Stoeckli, P. Robin Hiesinger, Bassem A. Hassan

## Abstract

Developmental variation in brain-wiring contributes to behavioural individuality^1,2^. However, how and when individualized wiring diagrams emerge and become stable during development remains largely unknown. Here, we explored axon targeting dynamics in individual brains using live-imaging of a developing *Drosophila* visual circuit and discovered that targeting choice is an algorithmic multi-step growth process with variable outcomes. Using optogenetics, we found that temporally restricted Notch lateral-inhibition defines a subset of neurons with a probabilistic potential to innervate distal targets. Next, axons from Notch^OFF^ neurons amplify into long actin-rich multi-fibre structures necessary for distal growth. A subset of these Notch^OFF^ neurons create distal targeting axons by stabilizing microtubule growth in one of their actin fibres. Amplified axons without tubulin-stabilized fibres retract, resulting in the stochastic selection of a different number of distal targeting axons in each brain. Pharmacological microtubule destabilization suffices to inhibit this targeting. We observed a similar axonal amplification-stabilization process in the developing chick spinal cord, suggesting a conserved mechanism. Finally, early microtubule patterns predict the adult brain-wiring of an individual in a target-independent manner prior to synapse formation^3,4^. Thus, we show that a temporal succession of genetically encoded stochastic processes explains the emergence of individual wiring variation.

**One-Sentence Summary:** The temporal succession of stochastic developmental processes explains the emergence of individual wiring variation.

---

Although brain-wiring patterns are similar between individuals, they are not identical. In all animals, including humans, structural variation between individuals sharing the same genetic and environmental background, and between the two hemispheres of the same brain, ensures behavioural individuality^5–7^ while maintaining the population around an optimal mean for niche exploitation^8,9^. *Drosophila melanogaster* is a powerful model to investigate stochastic variation^10^, individual behaviour^11,12^ and the relationship between the two^2,12–15^. Asymmetry in the *Drosophila* central nervous system has an impact on individual long-term memory^13^. Structural variations in a group of higher order neurons, called Dorsal Cluster Neurons (DCNs)/LC_14_ in the *Drosophila* visual system lead to individuality in visual response behaviour^2^. These neurons are divided in two sub-types: the lobula targeting DCNs (L-DCNs) and the medulla targeting DCNs (M-DCNs) **(Figure 1A, B)**. While we previously showed that within a DCN lineage, sub-fates are established post-mitotically^1^, it is unclear when, where and how during development DCN neurons become irreversibly committed to their sub-fate as L- or M-DCNs. The distinction between these two subtypes requires Notch signalling. However, while Notch activation prevents the innervation of the medulla in adult brains, *Notch* loss of function does not deterministically convert an immature DCN into a M-DCN. Rather, Notch^OFF^ status only increases the probability of innervating the medulla from ∼25% to ∼60%^1^. Thus, Notch signalling is a necessary but not sufficient step in DCN wiring specificity indicating that at least one further selection step is required to generate the final adult wiring pattern. What the potential second selection step is, how it relates to Notch activity, and how and when it contributes to the stabilization of axon targeting during development is unknown.

**Fig. 1.**
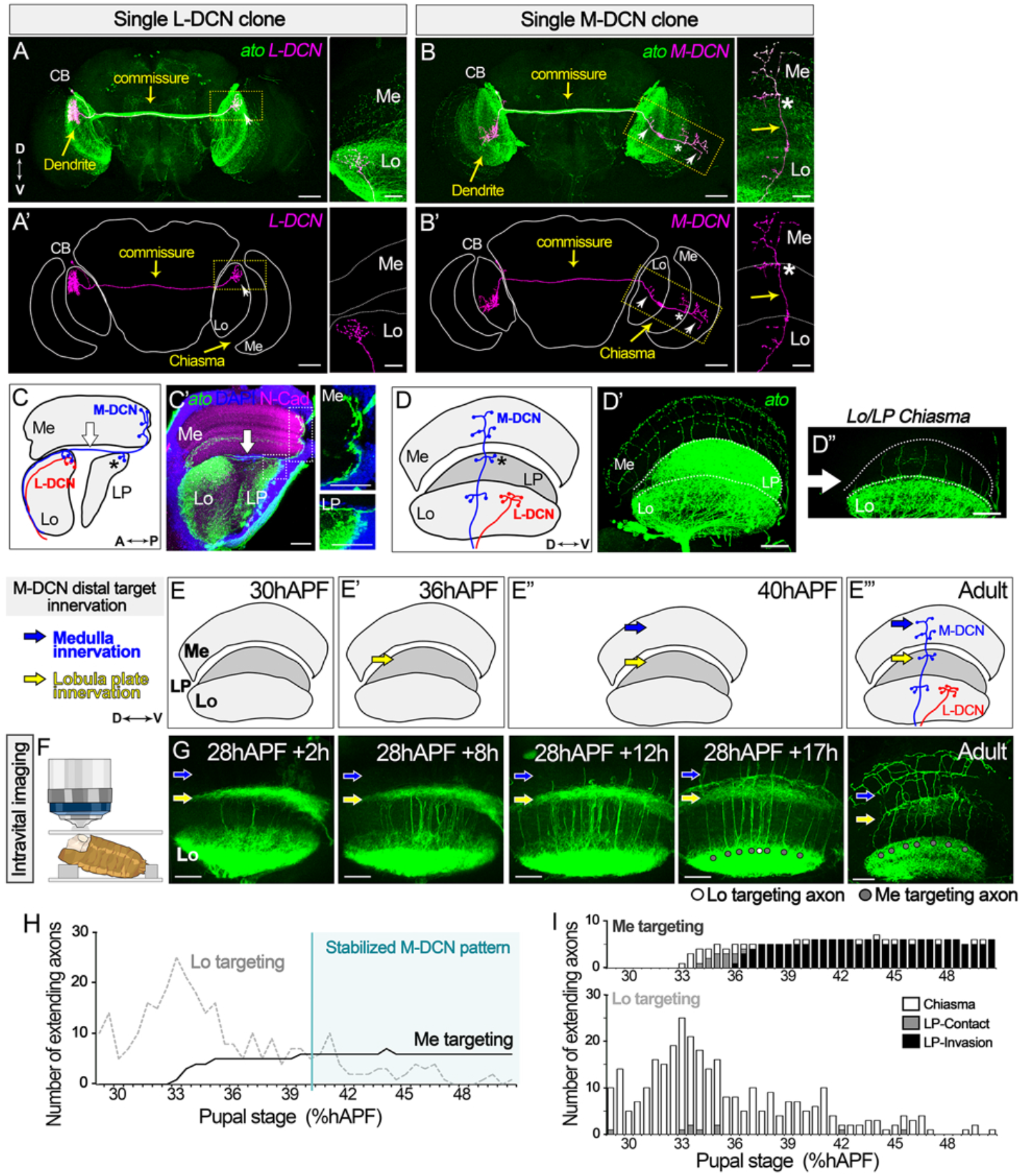
DCN Target choice precedes distal target innervation. **(A–B)** Single cell clones (magenta) of (A, A’) lobula targeting DCN (L-DCN) and (B, B’) medulla targeting DCN (M-DCN). White star: Lobula plate (LP). Arrowhead: lobula (Lo) and medulla (Me) innervations. White arrow: optic chiasma between the Lo and LP. Yellow arrow (High magnification): Chiasma. **(C-D)** Adult DCN innervation path in the optic lobe from (C) Dorsal and (D) Posterior view. (C’) M-DCN axons cross the chiasma in one plane (White arrow) to innervate the LP (black star) an Me. (D’-D”) M-DCN axons are organized in a fan-like shape along the Dorso-Ventral axis. Green: DCNs (CD4-tdGFP). Neuropile, N-Cadherin (N-Cad, magenta). Cell bodies (DAPI, blue). White arrow: Chiasma. **(E-E’’’)** Schematic of the sequential DCN target innervation: lobula (Lo), lobula plate (LP, yellow arrow) and medulla (Me, blue arrow). **(F)** Intravital imaging settings of the intact pupae. **(G)** Intravital live imaging of DCN axon development and the final adult pattern. Green: DCNs (CD4-tdGFP). Medulla targeting structures (gray circle) lobula targeting structures (white circle) **(H)** Number of lobula targeting and medulla targeting axonal structures through time from live imaging data. (**I**) Length of lobula and medulla targeting axonal structures through time from live imaging data. Scale bars (A-B):50µm and 20 µm; (C’)20µm; (D’)30µm; (G)20µm.

## Axon targeting decision occurs prior distal target innervation

L- and M-DCNs are defined by their final axonal targeting area. Each DCN projects a single axon towards the contralateral optic lobe to target two alternative regions^1,16–18^. Within the optic lobe, L-DCN axons terminate in the anterior lobula **(Figure 1A, C-D)**, while M-DCN axons additionally cross the optic chiasma between the lobula (Lo) and lobula plate (LP) **(Figure 1B-D”)** to innervate posterior-lateral LP targets and then terminally project in several layers of the posterior medulla **(Figure 1C-C’)**^19^. All DCNs are generated from a common progenitor^16,17^. L- and M-DCN sub-fates emerge at least in part through a cell-based competition from a pool of immature DCNs, with a minority of DCNs targeting the medulla^1^. We sought to understand when, where and how this post-mitotic sub-fate choice is made during development.

Two different strategies^20^ could explain medulla innervation by a DCN axon. In the case of a target-dependent mechanism, DCN axons would first reach distal targets and then retract if they do not find their medulla post-synaptic partners. In a target-independent mechanism, medulla targeting commitment would occur prior to axons reaching the medulla. In order to discriminate between these two models, we genetically tagged DCNs^16^ with two copies of the bright CD4-tdGFP^21^ and live-imaged the development of DCN axons using intravital 2-photon microscopy in intact animals^22^ **(Figure 1F)**. We first determined the developmental sequence of the DCN target innervation starting from the end of lobula innervation at 30 hours after pupa formation (hAPF) onward **(Figure 1E, G; Figure S1A)**. From 36hAPF, DCN axons reach the intermediate target: the lobula plate (LP), to finally commence medulla innervation around 40hAPF **(Figure 1E-E”)**. We found that the future M-DCN pattern became discernible at 44-50hAPF, suggesting the medulla targeting decision occurred before this time point **(Figure 1H, S1B-C; Movie S1)**. We took advantage of imaging intact animals to identify the fate of developing axons by comparing this stable developmental stage (44-50hAPF) with the final adult pattern and backtracking DCN axon behaviour within the lobula/lobula plate chiasma throughout development in individual flies **(Figure 1G, 1H, Figure S1B-D)**. We found that future medulla targeting axons extended to the lobula plate and medulla, and never retracted to the lobula. On the other hand, future lobula targeting axons extended into in the chiasma but did not innervate the lobula plate nor the medulla **(Figure 1H-I, S1D-E, S1G)**. Finally, by tracking medulla targeting and lobula targeting axons, we found that the final number of medulla targeting axons in the chiasma area in each individual is reached before medulla innervation (40hAPF) **(Figure 1H, S1D)**. Thus, DCN axon targeting decision is target independent and occurs prior to lobula plate and medulla innervation.

### Transient axon amplification is necessary but not sufficient for distal targeting

We then asked what – if anything – distinguishes future L-DCN from M-DCN axons during development. We found DCN axons form two types of dynamic developmental structures during their extension in the chiasma **(Figure 2A-B; Movie S2)**: single fibres that grow into the chiasm and retract back to the lobula **(Figure 2A-B, S1E, S1G)**, and multi-fibre structures that eventually resolve into single fibres **(Figure 2B, 2E, S1E-F)**. While some multi-fibre structures extend in the chiasma and then retract to the lobula **(Figure 2A, S1G)**, others lead to medulla targeting axons **(Figure 2A, 2C, S1E, S1G; Movie S2-S3)**. The complexity of multi-fibre structures is reduced through development **(Figure S2E)** with the stabilization of one (82%) or occasionally two (18%) fibres as the future medulla-targeting axon **(Figure 2A, 2C-D, S1E-F, S1H)**. This suggests that while a single fibre is not sufficient, the formation of multi-fibre structures is a necessary but insufficient prerequisite of medulla targeting **(Figure 2E)**.

**Fig. 2.**
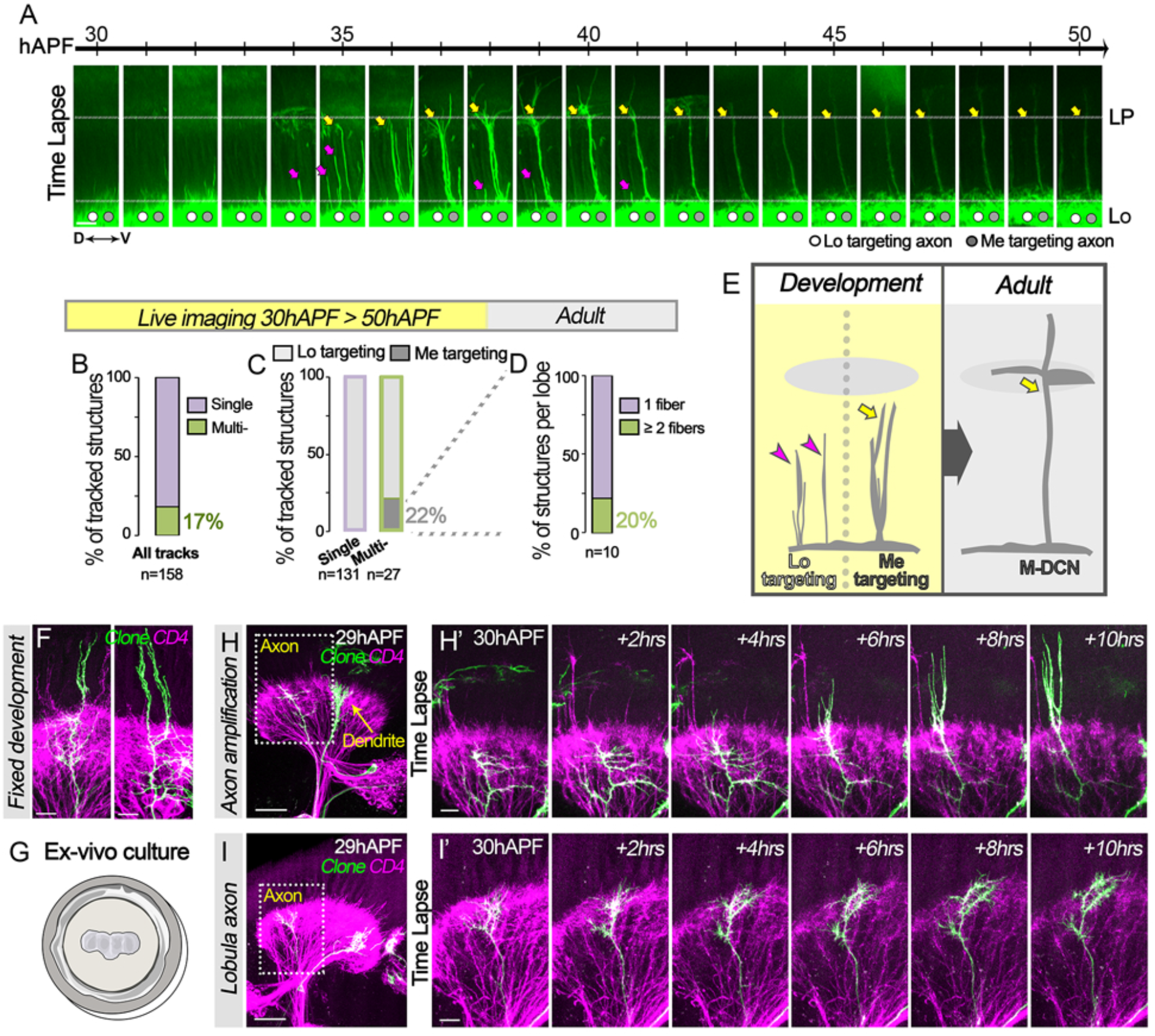
Single DCN axons amplify to form multi-fibre structures that are necessary but not sufficient for medulla targeting. **(A)** Intravital Imaging of developing DCN axons extending in the chiasma from the lobula (Lo), toward the lobula plate (LP). Medulla targeting structures (gray circle, Yellow arrow) lobula targeting structures (white circle, magenta arrow). Green: DCNs (CD4-tdGFP). (**B-D)** Single and multi-fibre structures were tracked over time during intravital live-imaging (B) and defined *a posteriori* as medulla or lobula targeting (C) based on the final adult pattern (D). **(E)** DCN multi fibre structures correlate with medulla targeting (Yellow arrow), while both multi- and single axonal fibre structures (magenta arrow) retract to the lobula. **(F)** Fixed imaging of a DCN amplifying axon clone (green). Magenta: CD4-tdGFP. **(G)** Schematic of the *ex-vivo* brain culture set up. **(H)** *Ex-vivo* live imaging of an amplifying DCN axon clone extending in the chiasma and **(I)** of a lobula exploring DCN axon clone. In green: DCN clone. Magenta: CD4-tdGFP. Scale bars: (A)10µm; (F)10µm; (H-I)30µm and 10µm.

To dissect the formation these multi-fibre structures at an higher spatio-temporal resolution, we generated GFP-labeled single-cell clones and performed live imaging in *ex-vivo* brain explants **(Figure 2G-I)**^23^. We found that individual axons can amplify to form these multi-fibre structures in the chiasma **(Figure 2F, 2H; Movie S4)**. We additionally found a population of DCN axons that locally explores the lobula, do not amplify in the chiasma, and sends unstable individual fibres into the chiasma **(Figure 2I; Movie S5)**. These data indicate that an early developmental step controls this axonal amplification in the chiasma and maintains a pool of axons in the lobula during development.

### Notch signalling temporally restricts axonal amplification

We investigated how these two axonal behaviours are regulated. Live imaging showed that two types of structures fail to reach the medulla: single fibres emanating out of the lobula, and a subset of multi-fibre structures that retract back from the chiasma **(Figure 2A-C, S1E-G; Movie S3)**. We previously showed that constitutive activation of Notch leads to an absence of medulla innervation in adult brains **(Figure 3A)**^1^. To determine how Notch activation impacts on DCN axon dynamics, we expressed an activated form of the Notch receptor in immature DCNs and tracked axon behaviour using intravital imaging. We found that while short individual fibres still extend into the chiasma from 30hAPF, no multi-fibre structures were observed, suggesting Notch activation prevents axon amplification and maintains axons in the lobula **(Figure 3B; Movie S6)**. We then asked whether Notch activation could also trigger axon retraction from the chiasma. We optogenetically^24^ activated Notch before and/or after the formation of multi-fibre structures in the chiasma **(Figure 3C-J)**. While activation throughout development and before axon extension in the chiasma prevented medulla innervation **(Figure 3C, F-G)**, activation during and after the formation of the multi-fibre structures did not **(Figure 3C, H-J)**. We then expressed an activated form of Notch receptor using a post-specification M-DCN GAL4 driver (VT037804) ^2^ and found that a Notch overactivation was not sufficient to revert a committed M-DCN into a lobula targeting neuron (**Figure S1I-J**). These data suggest that when DCN axons innervate the lobula from 0hAPF to 30hAPF, an early developmental step through Notch activation maintains a pool of axons in the lobula, while only Notch^OFF^ DCNs extend and amplify in the chiasma.

**Fig. 3.**
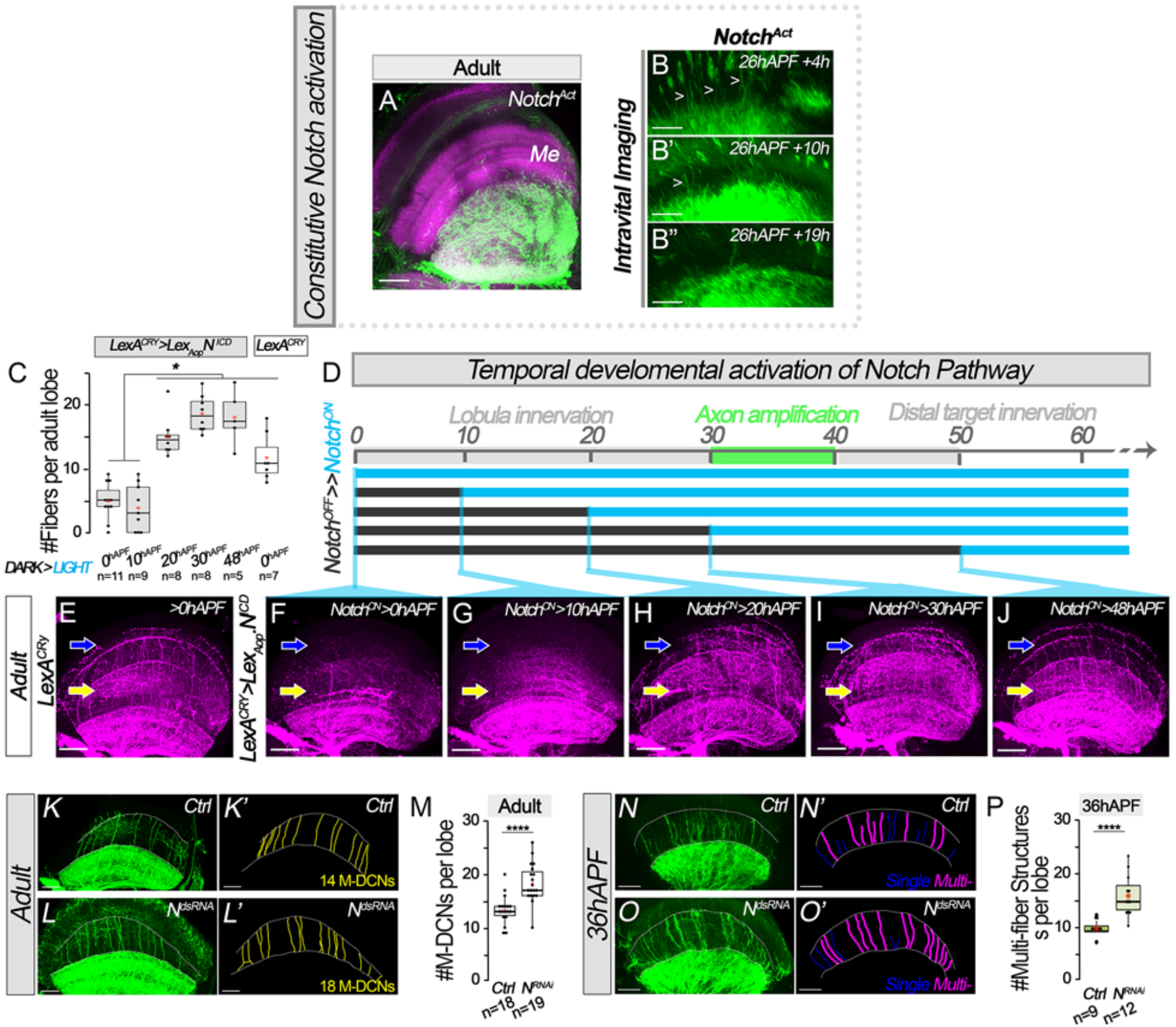
Notch signalling temporally restricts axonal amplification. **(A)** Constitutive activation of Notch receptor (Notch^Act^) during DCN development leads to an absence of medulla (Me) innervation in adult. Green: CD4-tdGFP. Neuropile, N-Cadherin (N-Cad, magenta). **(B)** Chiasma view during intravital live imaging of developing DCNs upon Notch overactivation (Notch^Act^). White arrow heads: Single axonal fibres. **(C-J)** Optogenetic temporal activation of Notch signalling pathway during DCN development. **(C)** Quantification of the number of axonal fibres per lobes upon a temporal activation of the Notch receptor. Mann-Whitney-Wilcoxon test: (*) p<0.05. Red star: Mean. **(D)** The split.LEXA-CRY system was activated through blue light exposure (Blue line) to trigger the expression of a constitutive active form of the Notch receptor (N^ICD^) in the DCN (F-G) before axon amplification (0hAPF or 10hAPF) or (H-J) during/after axon amplification (30hAPF or 48hAPF) in the chiasma..**(E)** As a control, a constant blue light exposure of individual without Notch intracellular domain (*N*^*ICD*^) transgene (*LexA*^*CRY*^) was used. Yellow arrow: chiasma. Blue arrow: medulla. **(K-P)** Tracking of the DCN single (blue) and multi-(magenta) axonal fibres in the chiasma (K-L) in adult and (N-O’) during development – 36hAPF upon *Notch*^*RNAi*^ compared to control (*Ctrl*) Green: CD4-tdGFP. **(P)** Quantification of the M-DCN axons per lobe in adult brains and **(P)** Quantification of the number of multi-fibre structures per lobe at 36hAPF in control *vs* Notch^RNAi^. Mann-Whitney-Wilcoxon test: (***) p<0.005. (****) p<0.001. Red star: Mean. Scale bars: (A)30µm; (B)15µm; (E-J)30µm; (K-l, N-O)20µm.

We then investigated the link between Notch^OFF^ status and axon amplification during development. *Notch* knockdown in DCNs led to more medulla innervation in the final adult DCN pattern^1^ **(Figure 3K-L’, M)** and a corresponding increase in multi-fibre structures at 36hAPF **(Figure 3N-O’; P)**, with no effect on the number of fibres per multi-fibre structures **(Figure S1K)**. All together our data so far show that lateral inhibition among DCNs^1^ defines two subpopulations: Notch^ON^ cells pre-committed to L-DCN fate and Notch^OFF^ cells, which amplify axons in the chiasma. However, only a subset of these Notch^OFF^ cells eventually becomes M-DCNs **(Figure 2C; S1G)**^1^, indicating the final selection of M-DCN fate requires further developmental steps.

### Microtubule growth selects future distal targeting axons

Although axon amplification correlates with DCN axon growth in the chiasma, our data indicate that stabilization of a single fibre within some of these structures is crucial for medulla innervation **(Figure S1G; Movie S3)**. To address this, we genetically labeled filamentous actin (F-Actin) using LifeAct^25^ and microtubules with the alpha-Tubulin84B subunit fused with GFP (Tubulin-GFP)^26^, specifically in DCNs. F-Actin is present in all fibres of an amplified axon **(Figure 4A)**. In contrast, Tubulin levels varied in time and space. An initially thick tubulin network at the base of the multi-fibre structure splits into two or more fibres, most with low intensity that grow, retract and eventually resorb **(Figure 4C-D, S2A-B; Movie S7-S8)**, and occasionally one with high intensity in a single fibre (Tub^GFP+^), which grows and never retracts (**Figure 4B-D, S2B; Movie S8-S9**). We observed a similar transient axonal amplification phenomenon during floor plate crossing of dI1 axons in chicken embryo spinal cord^27^. While crossing the midline, dI1 axons transiently split into two branches; one branch later retracts, while the other stabilizes and grows **(Movie S10)**. Live imaging of Tubulin dynamics in *ex-vivo* intact spinal cords revealed that a microtubule network splits and invades both branches. Tubulin then gradually disappears from one branch prior its full retraction, accumulates and stabilizes in the other branch that continues to grow (**Figure S2C; Movie S11**). Thus, a similar tubulin-based axonal fibre selection process also underlies floor plate crossing with the growth and stabilization of one single axonal fibre.

**Fig. 4.**
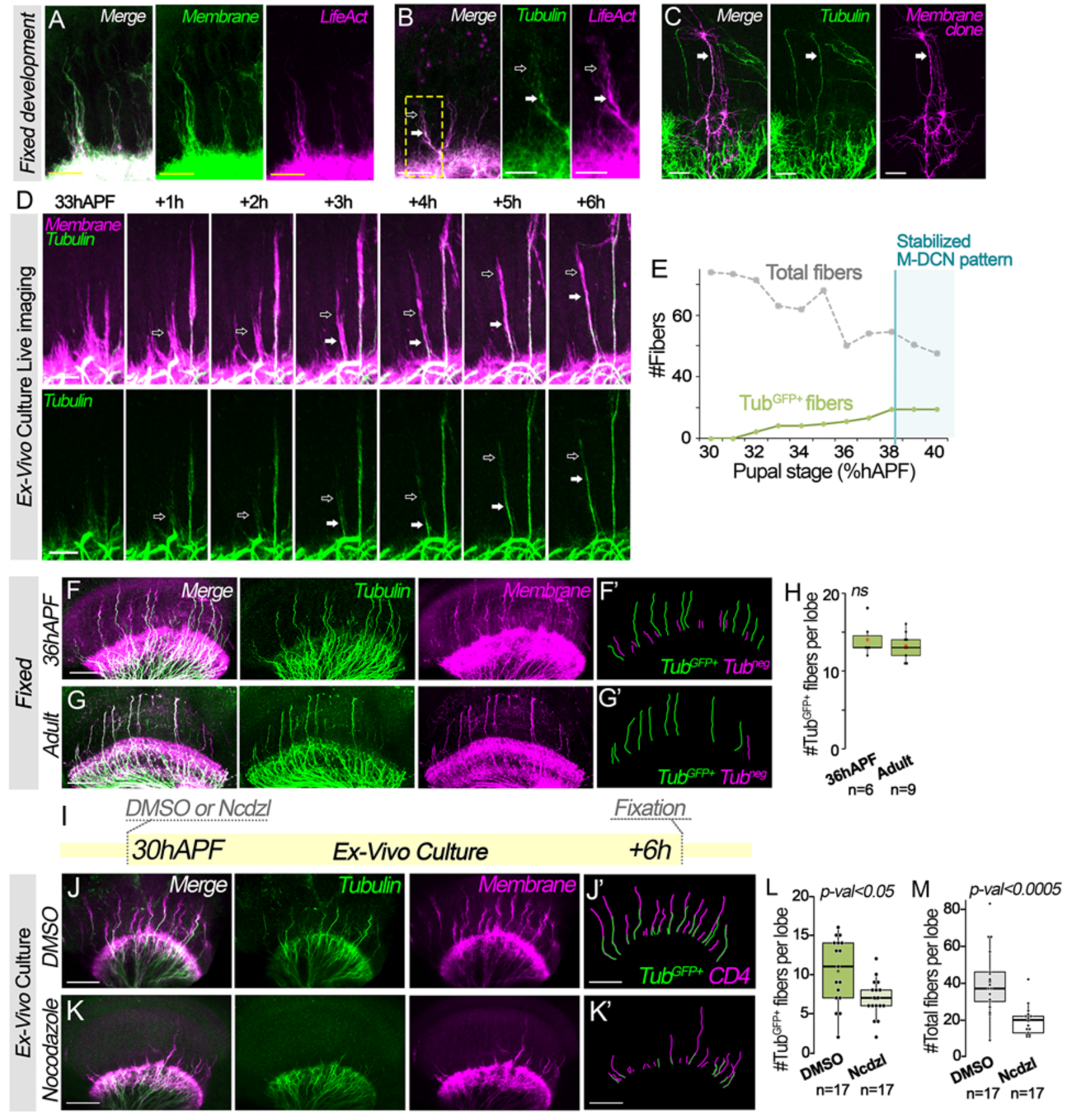
Microtubule growth selects future M-DCN axons. **(A-C)** DCN amplifying axons in fixed developing brains **(A)** F-Actin (LifeAct, magenta) is present in all fibres of a DCN amplifying axon. Membrane (CD4-tdGFP, green). **(B)** An amplifying axon contains two intensity levels of microtubules (green): a high amount of microtubule (white arrow) in one fibre, while the others have low amount of microtubule (black arrow). Actin: in magenta. **(C)** Single cell clone of an amplifying DCN axon (in magenta). Tubulin filaments (in green) innervate one of fibre of the amplifying axon (white arrow). **(D)** *Ex-vivo* brain culture live imaging of extending DCN amplifying axon shows microtubule fibres (white arrow) gradually extends and does not retract in the chiasma. Dynamic fibres with low amount of tubulin (black arrow) precede the microtubule fibre. Green: Tubulin, magenta: membrane. **(E)** Quantification of the total number of axonal fibres and tubulin fibres (Tub^GFP+^) through time. **(F-H)** Quantification of microtubule fibres (in green, Tubulin) in the chiasma at 36hAPF (F-F’, H) and in adult (G-G’, H) shows a similar number of microtubule fibres. **(I-N)** Microtubule destabilization from 30hAFP to 36hhAPF reduces axon extension. Green: Tubulin, magenta: membrane (CD4-tdGFP). **(L)** Quantification of the number of tubulin fibres and **(M)** the total number of fibres per lobe in the chiasma in control (DMSO) vs Nocodazole treatment (Ncdzl). Mann-Whitney-Wilcoxon test: (***) p<0.005. Red star: Mean.

Our analysis of microtubule dynamics during DCN outgrowth showed that some multi-fibre structures contain a Tubulin-rich fibre (Tub^GFP+^) that never retracts. We thus asked whether this one fibre is selected to become the future M-DCN axon. Using live imaging, we found that the number of fibres containing stable Tubulin gradually increases through development (**Figure 4D-E)** until ∼36hAPF when it stabilizes, while the total number of fibres decreases due to resorption back to the lobula (**Figure S2D**). The number of these stable Tub^GFP+^ fibres at ∼36hAPF precisely predicts the number of adult medulla targeting axons in each individual (**Figure 4F-H, S2E**), well before medulla innervation and synaptogenesis (50hAPF)^3,4^. We tested the idea that dynamic microtubule stabilization prior to ∼36APF is necessary for medulla targeting by treating *ex-vivo* cultured brains with Nocodazole between 30hAPF and 36hAPF **(Figure 4I-M)**. This led to a decrease in both Tub^GFP+^ fibres **(Figure 4L)** as well as the total number of fibres in the chiasma compared to controls **(Figure 4M, S3A-G; Movie S12-S13)**. In contrast, Nocodazole treatment at 48hAPF after the stabilization of the final pattern did not affect the number of medulla targeting axons, suggesting that microtubule stabilization is irreversible after this point **(Figure S3H-M**).

## Discussion

Our work addresses the question of the developmental origin of structural variation in adult brain wiring and shows that they emerge from a series of dynamic stochastic developmental processes. Within the fly visual system, the Dorsal Cluster Neuron (DCN/LC14) wiring diagram varies between the left and right hemispheres and among isogenic flies leading to individualized object response behaviour^2^. This variation relies on a differential innervation of proximal (lobula) versus distal (medulla) optic lobe neuropils by DCN axons. We previously showed that the probability of a DCN axon to target the medulla involves Notch cell-based competition between post-mitotic immature DCNs and is not instructed by lineage inherited factors^1^. Our current work highlights two successive stochastic developmental decision steps distinct in space and time, through which DCNs progressively lose their potential to target the medulla. First, while DCN axons are still in the lobula, the stochastic activation of Notch (Notch^ON^) maintains the majority of axons in there lobula, while Notch^OFF^ status permits formation of transient amplifying structures in the chiasma. Secondly, in a Notch-independent step, microtubules stabilize an individual fibre, or two in 20% of cases, within some of the Notch^OFF^ amplified axon, while the remaining fibres resorb. This leads to medulla innervation by single axons of small subset of DCNs, that thus become M-DCNs. Importantly, the statistical distribution of 0 to 2 selected axons from a given neuron underscores the probabilistic nature of the underlying selection process.

While Notch activation is sufficient to instruct the L-DCN fate, hence preventing the formation of M-DCNs, it did not induce the retraction of an amplifying axon, nor reverse a M-DCN committed axon. Through the course of axon development, a plethora of signals modulates the intrinsic states of an immature neuron until it reaches its final state^28^. For DCNs in particular, JNK signalling acts downstream of Notch to impact DCN medulla innervation potential^1^ and is additionally modulated by Wnt and FGF signalling^18^. We imagine that while a DCN axon grows in an external space of potential targets, it also navigates its intrinsic temporal states. Therefore, the spatio-temporally sequential activation of various locally derived signals, like Wnt and FGF among many others, would make a DCN non-responsive to further Notch activation and thus explain why specific signals can only act in specific temporal windows.

During the second phase of medulla targeting, we showed Notch^OFF^ DCN axons amplify into several long parallel actin-rich fibres generated from single cells. The formation of such axonal structures is not restricted to the DCN^27,29–31^. Mouse retinal ganglion cell axons display similar elongated growth cone (40µm) when navigating along the optic nerve^31^. Growth cones of dI1 neuron in the embryonic chicken spinal cord transiently amplify while crossing the floor plate (FP)^27^. This conserved growth cone behaviour may be induced by a change in substrate, when growth cones migrate from a dense neuropile to a sparser environment organized in “open channels”^32,33^. This radical change in morphology may also reflect a bet-hedging strategy as a growth cone crosses a non-adhesive environment, whereby the extension of multiple long parallel filopodia in the same direction would increase the probability of at least one fibre reaching the more adhesive target^34^. Similar to *Drosophila* DCNs, one fibre of the chick embryo spinal cord splitting axon is selected through microtubule stabilization, suggesting that microtubule-based axon amplification and stabilization is a conserved targeting mechanism.

Overall, our work supports a model **(Figure 5)** whereby the final innervation pattern, and thus the stabilization of a terminal fate of a neuron, emerges gradually in each brain through a process of genetically encoded algorithmic growth where the output of each step provides the input for a subsequent, but mechanistically independent ^35^. Consistent with this concept, early acting Notch activity defines the initial choice between L-DCNs and potential future M-DCNs, but is no longer required for the final selection of M-DCN axons. Conversely, late destabilization of tubulin reduces final medulla innervation, but does not prevent initial Notch-dependent entry into the chiasm. Perhaps most remarkably, we find that the final innervation pattern is variable across individual brains, cannot be precisely predicted from the mechanism, and must be actually observed until it is stabilized, consistent with the most salient prediction of the concept of wiring by algorithmic growth^36^.

**Fig. 5.**
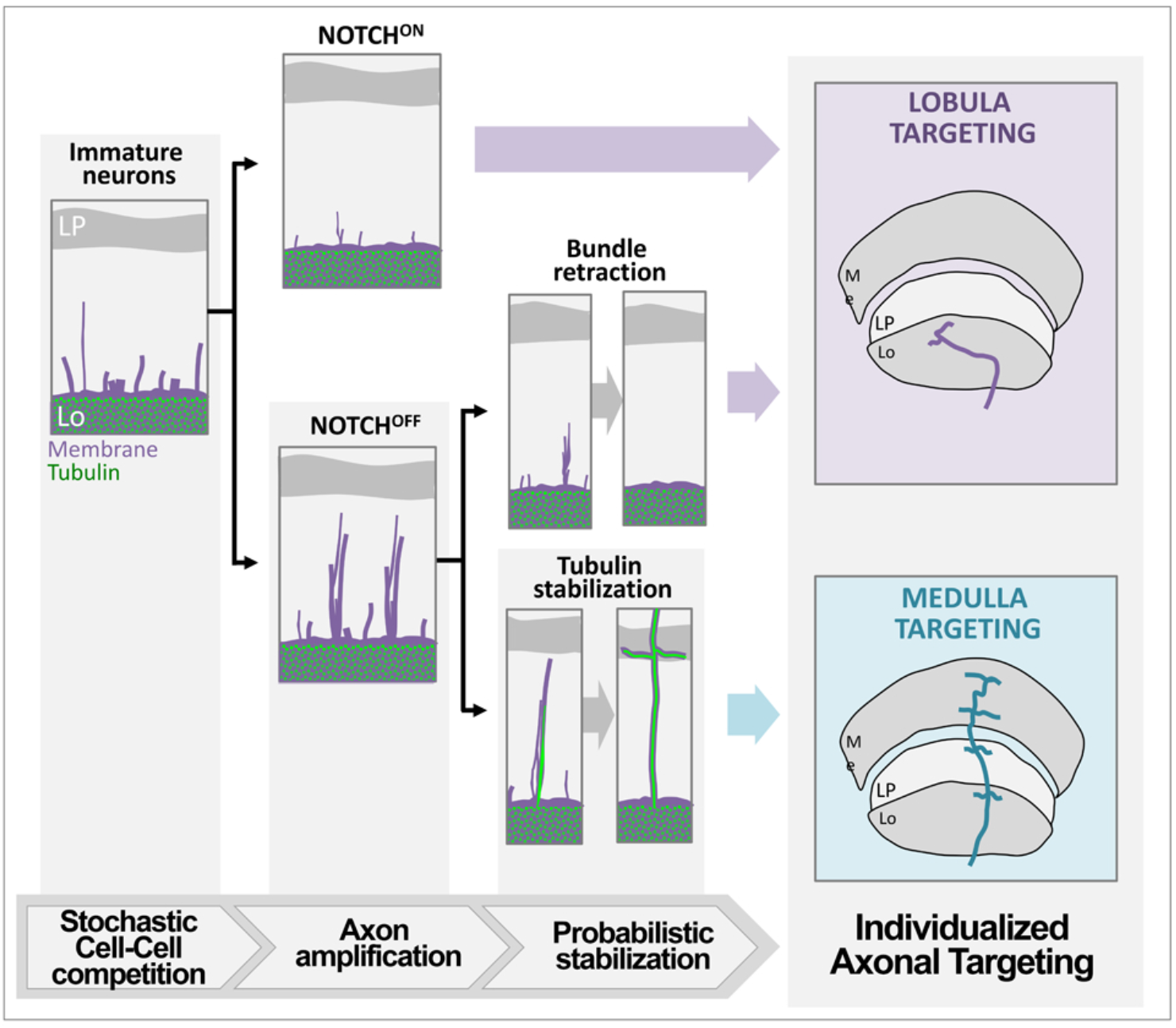
Model. A developmental succession of probabilistic steps creates individualized axonal wiring patterns. First, Notch lateral inhibition between immature post-mitotic DCNs defines which axons stay in the lobula (NOTCH^ON^). The second selection step relies on the capacity for the remaining Notch^OFF^ DCNs to amplify and form multi-fibre structures in the chiasma. Finally, a stochastic microtubule-dependent process selects one fibre from an amplifying axon as the future M-DCN axon shaft.

## Material and Methods

### Fly stocks and experiments

For all experiments, pupae were aged at 25°C and female *Drosophila melanogaster* were selected at 0hAPF (white pupae) based on the absence of male gonads. The following fly stocks were used: *ato*.*14A-GAL4* (*1*), *UAS-CD4-tdGFP* on 2^nd^ and 3^rd^ chromosomes (BDSC#35839 and BDSC#35836) (*2*), *UAS-CD4-tdTomato* (III) (BDSC#35837) (*2*), *pBPhsFlp2::PEST* (X) (*3*), *UAS-FRT-STOP-FRT*.*CD4-tdGFP, UAS-Notch*^*dsRNA*^ (BDSC#7077), *UAS-Notch*^*intra*.*GS*^ (BDSC#52008) (*4*), *lexAop-Notch*^*intra*^ (*5*), *UAS-CIBN::LexA-mcherry-p65::CRY2* (Gift from Kravitz lab)(*6*), *UAS-LifeAct*.*RFP* (III) (BDSC#58362), *UAS-Lifeact-Ruby* (II) (BDSC#35545), *UASp-GFPS65C-alphaTub84B* on 2nd and 3rd chromosomes (BDSC#7374 and BDSC#7373) (*7*), *pBPhsFlp2::PEST;;10UAS-HA_V5_FLAG* (*3*).

### Genotypes

The list of genotypes for all figures and supplementary material. <u>Figure 1</u>

**Fig. 1 A-B:** *pBPhsFlp2::PEST in attP3/w; +/+; ato*.*14A*.*GAL4, UAS-CD4*.*td-GFP/ pJFRC201-10XUAS-FRT>STOP>FRT-myr::smGFP-HA in VK0005, pJFRC240-10XUASFRT>STOP>FRT-myr::smGFP-V5-THS-10XUAS-FRT>STOP>FRT-myr::smGFP-FLAG*

**Fig. 1 D-I:** *w/w* ; *+/+; ato*.*14A*.*GAL4, UAS-CD4*.*td-GFP*

<u>Figure 2</u>

**Fig. 2 A-D:** *w/w* ; *+/+; ato*.*14A*.*GAL4, UAS-CD4*.*td-GFP*

**Fig. 2F:** *pBPhsFlp2::PEST in attP3/w; +/+; ato*.*14A*.*GAL4, UAS-CD4*.*td-GFP/ pJFRC201-10XUAS-FRT>STOP>FRT-myr::smGFP-HA in VK0005, pJFRC240-10XUASFRT>STOP>FRT-myr::smGFP-V5-THS-10XUAS-FRT>STOP>FRT-myr::smGFP-FLAG*

**Fig. 2 H-I:** *pBPhsFlp2::PEST in attP3/w; UAS-FRT>STOP>FRT-CD4*.*td-GFP/+* ; *ato*.*14A*.*GAL4, UAS-CD4*.*td-Tomato/+*

<u>Figure 3</u>

**Fig. 3A-B:** *w/w; UAS-Notch*^*intra*.*GS*^*/+; ato*.*14A*.*GAL4, UAS-CD4*.*td-GFP /+*

**Fig. 3C, F-J:** *w/w; LexAop-Notch*^*intra*^ */+; ato*.*14A*.*GAL4, UAS-CD4*.*td-Tomato, UAS-CIBN::LexA-mcherry-p65::CRY2/+*

**Fig. 3C, E:** *w/w; + /+; ato*.*14A*.*GAL4, UAS-CD4*.*td-Tomato, UAS-CIBN::LexA-mcherry-p65::CRY2/+*

**Fig. 3 K, M, N, P:** *w/w* ; *+/+; ato*.*14A*.*GAL4, UAS-CD4*.*td-GFP / +*

**Fig. 3 L, M, O, P:** *w/w* ; *+/+; ato*.*14A*.*GAL4, UAS-CD4*.*td-GFP / UAS-Notch*^*dsRNA#7077*^

<u>Figure 4</u>

**Fig. 4A:** *w/w; UAS-Lifeact-Ruby/+* ; *ato*.*14A*.*GAL4, UAS-CD4*.*td-GFP /+*

**Fig. 4B:** *w/w* ; *UASp-GFPS65C-alphaTub84B /+* ; *ato*.*14A*.*GAL4/ UAS-LifeAct*.*RFP*

**Fig. 4C:** *pBPhsFlp2::PEST in attP3/w* ; *UASp-GFPS65C-alphaTub84B /+* ; *ato*.*14A*.*GAL4/ pJFRC201-10XUAS-FRT>STOP>FRT-myr::smGFP-HA, pJFRC240-10XUASFRT>STOP>FRT-myr::smGFP-V5-THS-10XUAS-FRT>STOP>FRT-myr::smGFP-FLAG*

**Fig. 4D-M:** *w/w* ; *UASp-GFPS65C-alphaTub84B /+* ; *ato*.*14A*.*GAL4, UAS-CD4*.*td-Tomato*

<u>Supplementary Figures:</u>

**Fig. S1A-H:** *w/w* ; *+/+; ato*.*14A*.*GAL4, UAS-CD4*.*td-GFP / +*

**Fig. S1I:** *w/w* ; *+/+; VT037804*.*GAL4, UAS-CD4*.*td-GFP / +*

**Fig. S1J:** *w/w* ; *+/+; VT037804*.*GAL4, UAS-CD4*.*td-GFP / UAS-Notch*^*intra*.*GS*^

**Fig. S1K:** *w/w* ; *+/+; ato*.*14A*.*GAL4, UAS-CD4*.*td-GFP / +*

*w/w* ; *+/+; ato*.*14*.*GAL4, UAS-CD4*.*td-GFP / UAS-Notch*^*dsRNA#7077*^

**Fig. S2A-B, S3A-M:** *w/w* ; *UASp-GFPS65C-alphaTub84B /+* ; *ato*.*14A*.*GAL4, UAS-CD4*.*td-Tomato*

<u>Movies:</u>

**Movie 1-2:** *w/w* ; *+/+; ato*.*14A*.*GAL4, UAS-CD4*.*td-GFP / +*

**Movie 3-4:** *pBPhsFlp2::PEST in attP3/w; UAS-FRT>STOP>FRT-CD4*.*td-GFP/+* ;

*ato*.*14A*.*GAL4, UAS-CD4*.*td-Tomato/+*

**Movie 5: Fig. 3G-H:** *w/w; UAS-Notch*^*intra*.*GS*^*/+; ato*.*14A*.*GAL4, UAS-CD4*.*td-GFP /+*

**Movie 6-7-8-13-14:** *w/w* ; *UASp-GFPS65C-alphaTub84B /+* ; *ato*.*14A*.*GAL4, UAS-CD4*.*td-Tomato*

### Clonal Induction

Multi-color flip out (MCFO) clone induction for fixed analysis [Fig. 1A-B; Fig. 2F] and CD4-td.GFP flip-out clone induction for live imaging [Fig. 2H-I]. White pupae were collected and heat-shocked for 15min at 37°C, and then grown until the indicated developmental age or adulthood.

### Temporal Optogenetic activation of Notch

We used the UAS-LEXA.CRY system (Chan et al. 2015) to control the expression of the Notch receptor intracellular domain (NICD). The split LEXA-p65 transcription factor (TF) is expressed in the DCN under the control of the ato14A.GAL4 driver and is inactive in dark. Upon Blue light exposure, the CIBN/CRY system reconstitutes a functional LEXA-p65 TF, which activates the transcription of NICD.

Fly crosses were kept in dark at 25°C. Females F1 white pupae (0hAPF) were collected under red light and raised in dark at 25°C until the indicated time point of blue light exposure. For the optogenetic induction, pupae were then transferred in a vial and put under constant blue light (Blue LED, tension 24V, from LED-motion Ref. 6302) until adulthood with the following parameters – *Photons per surface unit per second*: Np = 3.5^e17^ photons.m^-2^.s^-1^. *Irradiance*: I = 1.5^e-12^ W.m^-2^. *Wavelength*: 470 nm.

### Immunostaining and fixed Imaging

An immunostaining was done for all the fixed data sets. Pupal and adult brains were dissected in PBS, fixed 20min in 3.7% paraformaldehyde (PFA). PBS. Washes were done in PBS, 0.4% Triton-X (PBST). Antibody incubations were performed in PBST, normal Donkey serum 1:200 (S30-100ML, Sigma-Aldrich), overnight at 4°C for primary antibodies and 3h at room temperature for secondary antibodies. Samples were mounted in Vectashield (Vector Laboratories, CA) and imaged using a 63X glycerol objective (NA=1.3) with a Leica TCS SP8-X white laser confocal microscope.

primary antibodies and 3h at room temperature for secondary antibodies. Samples were mounted in Vectashield (Vector Laboratories, CA) and imaged using a 63X glycerol objective (NA=1.3) with a Leica TCS SP8-X white laser confocal microscope.

<u>The following antibodies were used</u>: anti-GFP (chicken, Abcam; ab13970, 1:500), anti-mCherry (rabbit, Abcam; #ab167453, 1:500) or anti-DsRed (rabbit, ClonTech; 1:300) to stain against Tomato, anti-NCadherin (rat monoclonal, Developmental Studies Hybridoma Bank; DN-Ex #8, 1:50), anti-Flag DYKDDDDK Epitope (rat monoclonal, Novusbio; NBP1-067-12 clone L5, 1:200), anti-HA (rabbit monoclonal, Cell Signalling; C29F4, 1:300), anti-V5 clone SV5-Pk4 (mouse monoclonal, BioRad; MCA1360, 1:500), Alexa 405, 488, 554, 647, (Donkey, Jackson ImmunoResearch Laboratories, 1:500).

### Drosophila Intravital mounting

Pupae expressing two copies of the *UAS-CD4*.*td-GFP* transgene under the control of the promoter *ato14A-GAL4* (*1*) were staged and aged at 25°C from 0 hours after puparium formation (hAPF) until the indicated hour for imaging. The cuticle surrounding the pupal head was then removed. Pupae were mounted by following the protocol described in Langen et al. 2015, to image the DCNs through the right eye. In order to compare the developing and adult DCN patterns from the same brain, pupae were then removed from the chamber after imaging and kept at 25°C until adulthood. The corresponding one-week-old adults were dissected, fixed and stained for further confocal imaging.

### Drosophila Brain culture

*Ex-vivo Drosophila* brain cultures were performed according to the methods described by Ozel et al. 2015(*9*). Pupal brains were dissected one hour before the imaging time point in Schneider’s Drosophila Medium. Two layers of 1×1 mm squares of 200 mm tape were used as spacers.

### Nocodazole treatment

Pupal brains were dissected in Schneider’s Drosophila Medium 30min before the indicated time point (30hAPF or 48hAPF) of drug treatment, and embedded at the indicated time point into 0.4% low-melting agarose in culture medium (see Ozel et al. 2015) *(8*) complemented with 10µg/mL [33µM] final concentration of Nocodazole (SML1665-1ML, Sigma) or DMSO (D8418-50ML, Sigma). Embedded brains were then kept at 25°C in DMSO or Nocodazole supplemented culture medium for 6hours. After treatment, brains were removed from the Low-melted Agarose, fixed and immunostained as described above. For Ex-vivo live brains in Low-melted agarose were mounted as the regular protocol with drug supplemented Culture medium and were imaged 30min-1h after the drugs were added.

### Live Imaging acquisitions (Drosophila)

All *Drosophila* Live imaging experiments were acquired on a Leica SP8 MP 2-photon microscope with a 40X IRAPO water objective (NA=1.1), a Chameleon Ti:Sapphire laser and Resonant scanner with a single excitation laser at 896nm for either one color imaging (GFP) or two color imaging (GFP/Tomato). The Nocodazole Live imaging experiments were acquired on a Ultima 2Pplus Bruker 2-photon microscope with a N20X-PFH - 20X Olympus XLUMPLFLN Objective (1.00 NA, 2.0 mm WD) and Resonant scanner, 896nm excitation laser.

### Image processing & analysis

During pupal development, the optic lobe undergoes a global rotation during the first half of the metamorphosis (*10*). Similarly, the Medulla rotates and is pulled by the Lamina from 25hAPF (*11, 12*). These rotations could be observed during our in-vivo Intravital live-imaging and were corrected using the ‘Image Alignement’ module from Imaris (Bitplane, Switzerland). To quantify the number of total axonal structures (single and multi-fibers), the number of axonal fibers, the percentage of multi-fiber structures and the number of fibers per axonal structure [fig. 1H, fig. S1C, fig. S1G-J, fig. 2D, fig. 3E, fig. 3F, fig. 3J, fig. 4E, fig. 4H, fig. 4L] for our fixed data set and each time point of our 4D data sets, axonal structures in the lobula/lobula plate chiasma were skeletonized using the ‘Filament Tracer’ module from Imaris (Bitplane, Switzerland). A structure was defined as “multi-fiber” when two or more fibers shared a least one contact point. To track axonal structures through time (fig. 1I, fig. S1D, fig. S1F, fig. S1F, fig. 2B, fig. 2C), which we called a “targeting event”, we used the ‘TrackingLineage’ module from Imaris (Bitplane, Switzerland). Statistical analyses were performed using the non-parametric unpaired two-samples Mann-Whitney-Wilcoxon test in R. A Kolmogorov–Smirnov test was additionally performed in R to compare distributions (fig. S1 H-J).

### In ovo electroporation

We used a plasmid encoding membrane-bound farnesylated td-Tomato (td-Tomato-F) under the control of the Math1 enhancer and the β-globin promoter (*13*) to specifically label dI1 commissural axons in the chicken spinal cord. The chicken tubulin-GFP sequence was subcloned from a pCAG tubulin-GFP plasmid into one with the Math1 enhancer and the β-globin promoter. pCAG tubulin GFP was a gift from Felicitas Proels & Martin Scaal (Addgene plasmid #66105 ; http://n2t.net/addgene:66105 ; RRID:Addgene_66105)(*14*). *In ovo* electroporation was performed as described previously in a video protocol (*15*). Briefly, plasmids were injected alone (td-Tomato-F plasmid) or together with the tubulin-GFP plasmid at a concentration of 700 ng/µl each in the central canal of the neural tube of stage HH17-18 chicken embryos (*16*). A final concentration of 0.01% (vol/vol) of Fast Green was added to the plasmid mix to trace injection site and volume of the mix. Plasmids were unilaterally electroporated, using a BTX ECM830 square-wave electroporator (five pulses at 25 V with 50 ms duration each). After electroporation, embryos were covered with sterile PBS and eggs were sealed with tape and incubated at 39°C for 26–30 h, until embryos reached stage HH22.

### Live imaging of intact chicken spinal cords

Dissection, mounting and imaging of intact spinal cords were performed as previously described (*13*). Intact spinal cords were dissected from HH22 embryos and embedded, with the ventral side down, in a drop (100 µl) of 0.5% low-melting agarose (FMC) containing a 6:7 ratio of spinal cord medium (MEM with Glutamax (Gibco) supplemented with 4 mg/ml Albumax (Gibco), 1 mM pyruvate (Sigma), 100 Units/ml Penicillin, and 100 µg/ml Streptomycin (both Gibco) in a 35-mm Ibidi m-Dish with glass bottom (Ibidi, #81158). Once the agarose polymerized, 200 µl of spinal cord medium were added to the drop and live imaging was started. Live imaging recordings were performed with an Olympus IX83 inverted microscope equipped with a spinning disk unit (CSU-X1 10’000 rpm, Yokogawa). Cultured spinal cords were kept at 37°C with 5% CO_2_ and 95% air in a PeCon cell vivo chamber. Temperature and CO_2_-levels were controlled by the cell vivo temperature controller and the CO_2_ controller units (PeCon). Spinal cords were incubated for at least 30 min before imaging was started. We acquired 35-40 planes (1.5 µm spacing) of 2 × 2 binned z-stack images every 5 or 10 min for 12-24 hr with a 40x silicon oil objective (UPLSAPO S40x/1.25, Olympus) and an Orca-Flash 4.0 camera (Hamamatsu) with the help of Olympus CellSens Dimension 2.2 software. Maximum projections of Z-stack videos were generated and modified using Fiji/ImageJ (*17*). Heat maps of tubulin-GFP intensities were generated with the “Royal” Lookup table and growth cone boundaries were traced using the wand tool in Fiji/ImageJ.

## Acknowledgments

The authors thank the Bloomington stock center (NIH P40OD018537), the Vienna Drosophila Resource Center, the Developmental Hybridoma Bank, Pr. Edward Giniger and Pr. Edward A. Kravitz and Dr. Amr Hasan for flies and reagents; Dr Beat Kunz and Tiziana Flego for excellent technical, as well as Liz Hellbruegge and Ryan Cook for technical assistance; Pr. C. Desplan, all members of the Hassan, Hiesinger, and Wernet labs for support, insightful discussions and valuable comments. Light microscopy was carried out at in the Lab of Pr. P. Robin Hiesinger at Freie Universität Berlin and at the ICM.Quant facility in Paris.

## Author contributions

M.A., B.A.H. and P.R.H. conceived the study, designed the experiments, and wrote the manuscript.

M.A., A.D. and S.D. conducted all experiments and data analysis.

E.S., P.R.H. and B.A.H. acquired funding.

## Data and materials availability

All data are available in the manuscript or the supplementary materials.

## Extended Data figures

**Fig. S1.**
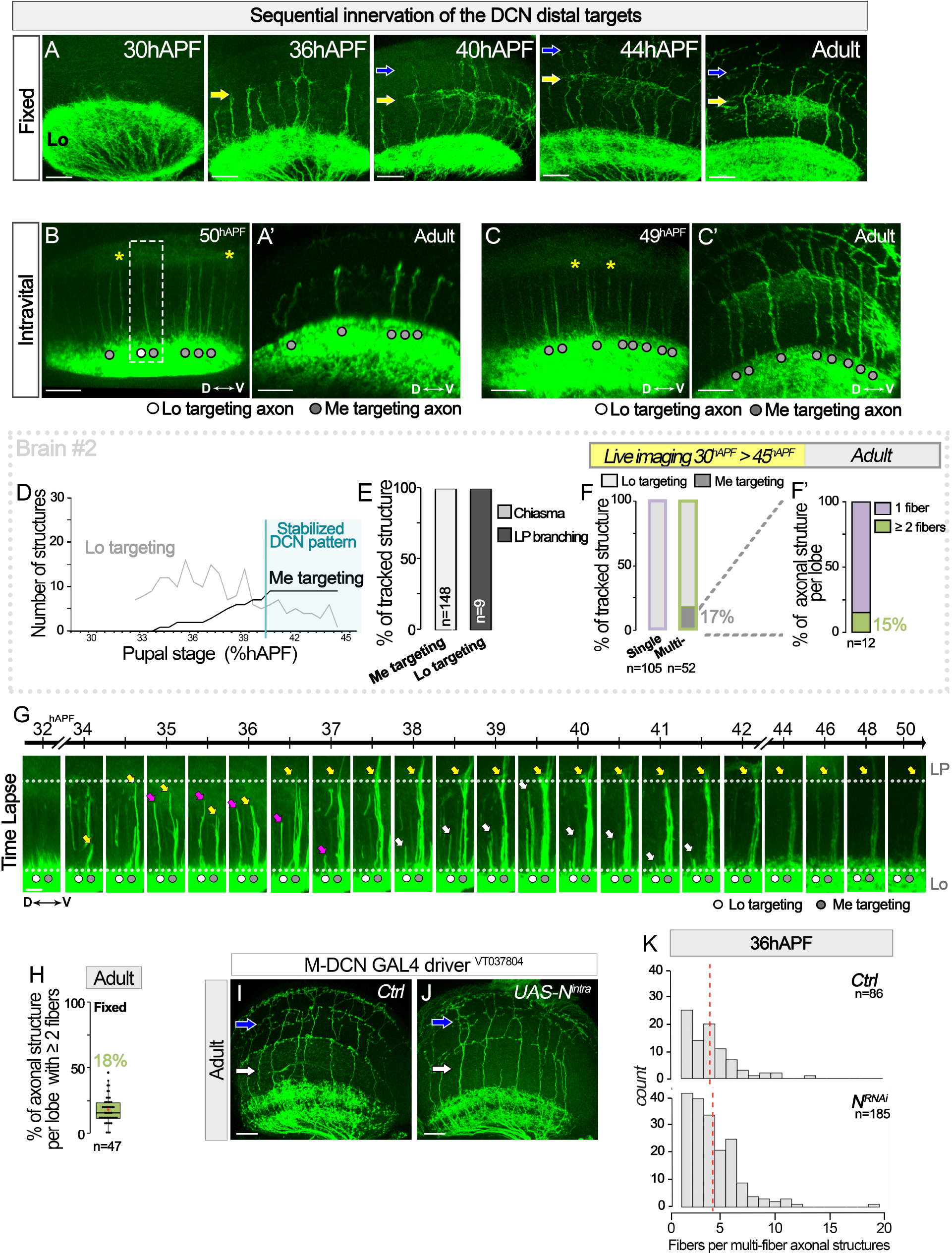
**(A)** Validation of the sequential DCN target innervation in fixed brain through development from 30 hour after pupation (hAPF) to adult. **(B-C)** Stabilized DCN pattern of brain developing from Figure 2A-D and their respective final adult pattern (B’-C’). In green: DCNs (CD4-tdGFP). Yellow stars: neuronal fibers that extend from the lobula (Lo) to the lobula plate (LP). **(D-F)** Single and multi fiber structures extending in the chiasma were tracked over time during Intravital live-imaging and defined a posteriori as medulla or lobula targeting (E) based on the final adult pattern (D) Number of medulla (Me) and lobula (Lo) targeting axonal structures through time from live imaging data. (E) Maximum length of Me- and Lo-targeting tracked axonal structures through (F) development and (F’) the final adult structure. **(G)** Intravital Imaging of developing DCN axons extending in the chiasma from the lobula (Lo), toward the lobula plate (LP) showing a multi-fiber structure targeting the medulla (Yellow arrow), a multi-fiber structure (white arrow) and a single fiber (magenta arrow) both retracting to the lobula. Medulla targeting structures (gray circle), lobula targeting structures (white circle). Green: DCNs (CD4-tdGFP). **(H)** Quantification of the percentage of multi fiber structures per lobe in adult fixed brains. Red star: Mean. **(I)** Notch overactivation (UAS-Nintra) in committed M-DCN during development did not prevent medulla targeting in adult. **(K)** Distribution of the number of fibers per multi fiber structures in control vs NotchRNA at 36hAPF (mean.ctrl= 4.06, mean.NRNAi= 4.33; Mann-Whitney-Wilcoxon test: p= 0.33; A Kolmogorov–Smirnov test: p= 0.91) Scale bars: (A-B, I-J)20µm; (G)10µm.

**Fig. S2.**
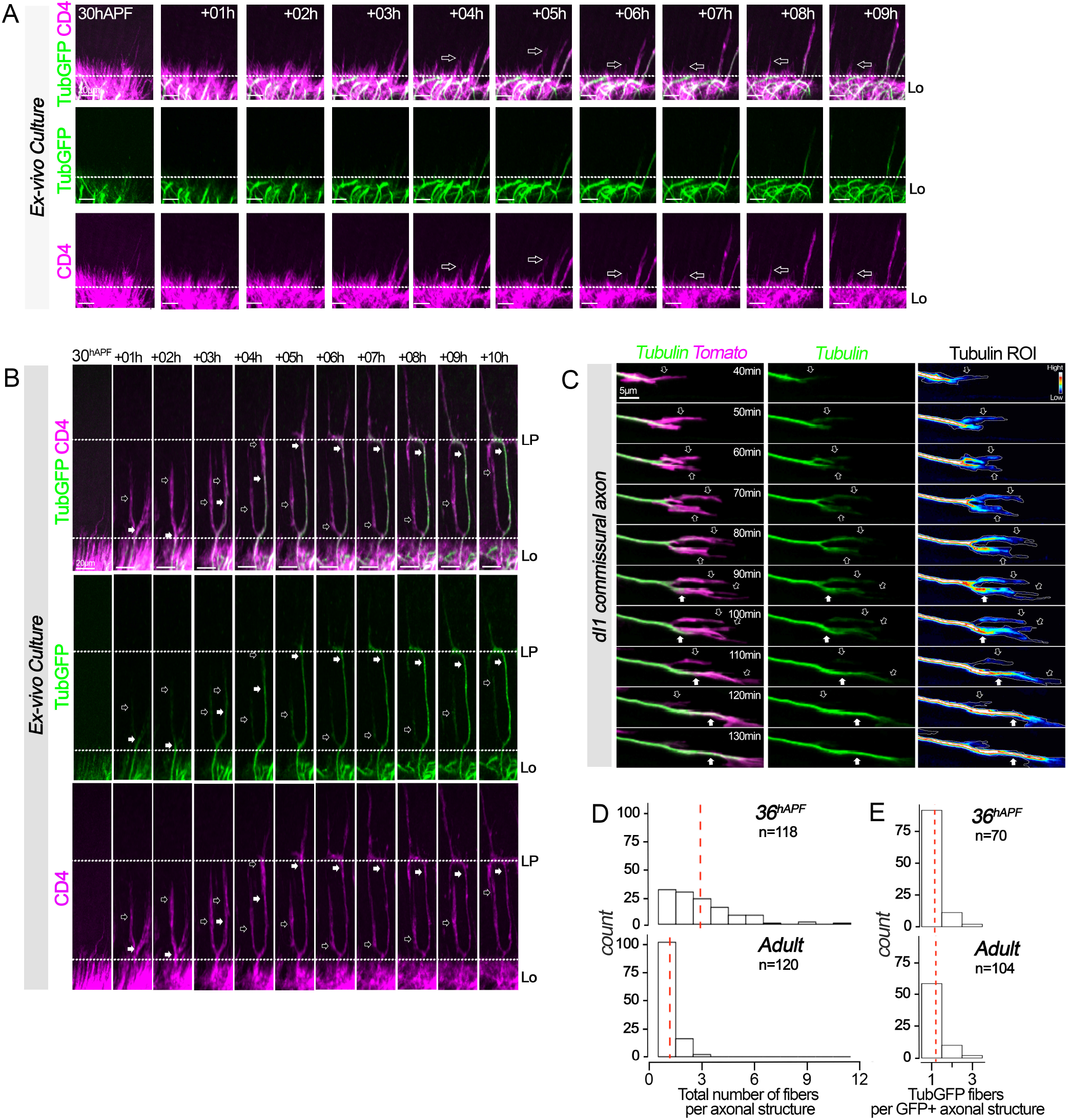
**(A-B)** *Ex-vivo* brain culture live imaging of (A) multi-fiber structures extending in the chiasma and then retracting to the lobula (Lo). (B) Low amount of Tubulin (black arrow) fluctuates between axonal fibers until it stabilizes one (white arrow). Green: Tubulin, magenta: membrane. **(C)** Live imaging of tubulin dynamics in transient dI1 axon growth cone splitting. Tubulin-GFP (in green) first localized to both branches (black arrowhead) at a low intensity, further stays and accumulates in the stabilized one (white arrowhead) and resorbs in the retracting one. Magenta: tdTomato-F. The heat map shows the tubulin-GFP signal in the growing axon (white line). Hi, high; Lo, low. **(D)** Distribution of the total number of fibers per axonal structure in the chiasma at 36hAPF and in adult brains (mean.36hAPF= 2.90, mean.Adult= 1.17; Mann-Whitney-Wilcoxon test: p<0.001; A Kolmogorov–Smirnov test: p<0.001). **(E)** Distribution of the number of Tubulin+ fibers (TubGFP) per axonal structures that contain TubGFP+ fibers in the chiasma at 36hAPF and in adult brains (mean.36hAPF= 1.20, mean.Adult= 1.14; Mann-Whitney-Wilcoxon test: p= 0.39; A Kolmogorov–Smirnov test: p= 1.00).

**Fig. S3.**
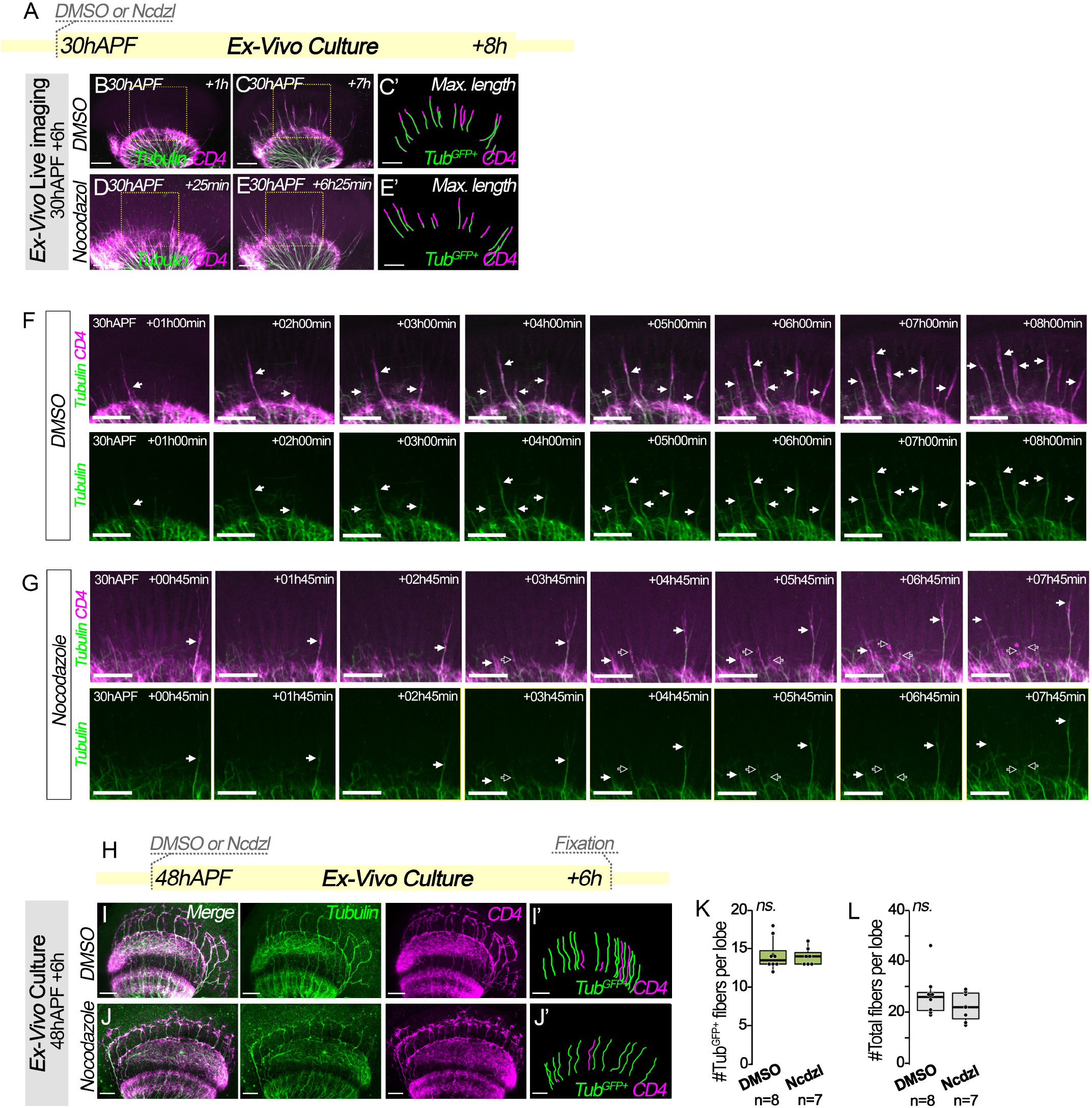
**(A-G)** Live imaging of *Ex-vivo* cultured brains treated with Nocodazole or DMSO during axon amplification. Green: Tubulin, magenta: membrane (CD4-tdGFP). High tubulin level (White arrow), Low tubulin level (Black arrow). **(H-M)** *Ex-vivo* cultured brains treated with Nocodazole (Ncdzl) or DMSO after M-DCN pattern stabilization from 48hAPF. **(K)** Quantification of the number of tubulin fibers and **(L)** the total number of fibers per lobe in the chiasma in control (DMSO) vs Nocodazole treatment (Ncdzl). Mann-Whitney-Wilcoxon test. Red star: Mean. Scale bars: (A-G)30µm; (I-J)20µm.

**Movie S1. Intravital imaging of DCN developing axon**. Related to Figure 1G. 21.5 hours time-lapse recording sequence (one stack every 20 min), starting from 30hAPF. White arrowhead: stabilized axonal structure. Green: CD4.tdGFP. Scale bar: 20µm.

**Movie S2. Intravital imaging of DCN developing axon in the Lo/LP chiasma**. Related to Figure 2A. 22.33 hours time-lapse recording sequence (one stack every 30 min), starting from 30hAPF. White arrowhead: stabilized axonal structure. Green: CD4.tdGFP. Scale bar: 10µm.

**Movie S3. Intravital imaging – Retracting DCN multi-fiber axonal structure in the Lo/LP chiasma**. Related to Figure S1G. 22.33 hours time-lapse recording sequence (one stack every 30 min), starting from 30hAPF. White arrowhead: stabilized axonal structure. Green: CD4.tdGFP. Scale bar: 10µm.

**Movie S4. *Ex-vivo* live imaging of DCN axon clone extending toward the chiasma with axonal amplification**. Related to Figure 2H. 14.33 hours time-lapse recording sequence (one stack every 20 min), starting from 29hAPF. Green: CD4.tdGFP clone. Magenta: CD4.tdTomato. Scale bar: 10µm.

**Movie S5. *Ex-vivo* live imaging of DCN axon clone exploring the lobula**. Related to Figure 2I. 10 hours time-lapse recording sequence (one stack every 6min 44s), starting from 29hAPF. Green: CD4.tdGFP clone. Magenta: CD4.tdTomato. Scale bar: 10µm.

**Movie S6. Intravital imaging of DCN developing axon in the chiasma upon a Notch singling pathway overactivation (*Notch***^***Act***^). Related to Figure 3B. 19hours time-lapse recording sequence (one stack every 15min), starting from 26hAPF. White arrowhead: single fiber structures. Green: CD4.tdGFP. Scale bar: 10µm.

**Movie S7. *Ex-vivo* live imaging of Tubulin dynamic in developing DCN axon in the chiasma**. Related to Figure 4D. 9hours 29min time-lapse recording sequence (one stack every 20 min), starting from 30hAPF. Green: tubulin-GFP. Magenta: CD4.tdTomato. Scale bar: 10µm.

**Movie S8. *Ex-vivo* live imaging of unstable multi-fiber structure from the chiasma to the lobula**. Related to Figure S2A. 9hours 29min time-lapse recording sequence (one stack every 20 min), starting from 30hAPF. Green: tubulin-GFP. Magenta: CD4.tdTomato. Scale bar: 20 µm.

**Movie S9. *Ex-vivo* live imaging of Tubulin dynamic in developing DCN axon in the chiasma**. Related to Figure S2B. 9hours 29min time-lapse recording sequence (one stack every 20 min), starting from 30hAPF. Green: tubulin-GFP. Magenta: CD4.tdTomato. High tubulin level (White arrow), Low tubulin level (Black arrow). Scale bar: 20µm.

**Movie S10. Live imaging of dI1 axon growth cone transient splitting during midline crossing in chicken embryo spinal cord**. 145-min time-lapse recording sequence (one stack every 5 min). Black arrows: the consecutive transient splitting events. White arrowhead: the stabilized branch. Black arrowheads: the retracting branch. Scale bar: 10µm.

**Movie S11 Live imaging of tubulin dynamics in dI1 axon growth cone transient splitting**. Related to Figure S2C. 240-min time-lapse recording sequence (one stack every 10 min). White arrows: transient growth cone. Black arrowheads: retracting branch. White arrowheads: stabilized branch. Green: tubulin-GFP. Magenta: tdTomato-F. The lower panel shows a heat map visualization of the tubulin-GFP signal. The white line shows the edge of the growing axon. Hi, high; Lo, low. Scale bar: 10µm.

**Movie S12. *Ex-vivo* live imaging of Tubulin dynamic in developing DCN axon in the chiasma – DMSO treatment from 30hAPF**. Related to Figures S3B-C, S3F. 8hours time-lapse recording sequence (one stack every 20 min). Green: tubulin-GFP. Magenta: CD4.tdTomato. White arrow: Stabilized Tubulin-rich axons. Scale bar: 30µm.

**Movie S13. *Ex-vivo* live imaging of Tubulin dynamic in developing DCN axon in the chiasma – Nocodazole treatment from 30hAPF**. Related to Figures S3D-E, S3G. 8hours time-lapse recording sequence (one stack every 20 min). Green: tubulin-GFP. Magenta: CD4.tdTomato. White arrow: Stabilized Tubulin-rich axons. Black arrow: Retracting axon. Scale bar: 30µm.

## References

1. Langen, M. et al. Mutual inhibition among postmitotic neurons regulates robustness of brain wiring in Drosophila. Elife (2013) doi:10.7554/eLife.00337.

2. Linneweber, G. A. et al. A neurodevelopmental origin of behavioral individuality in the Drosophila visual system. Science (80-.) . 367, 1112–1119 (2020).

3. Chen, Y. et al. Cell-type-specific labeling of synapses in vivo through synaptic tagging with recombination. Neuron 81, (2014).

4. Dutta, Suchetana ; Linneweber, Gerit Arne ; Andriatsilavo, Maheva ; Hiesinger, Peter Robin and Hassan, B. A Critical Developmental Interval of Coupling Axon Branching to Synaptic Degradation During Neural Circuit Formation. bioRxiv (2022).

5. Goodman, C. S. Isogenic grasshoppers: Genetic variability in the morphology of identified neurons. J. Comp. Neurol. (1978) doi:10.1002/cne.901820408.

6. Mohr, A., Weisbrod, M., Schellinger, P. & Knauth, M. The similarity of brain morphology in healthy monozygotic twins. Cogn. Brain Res. (2004) doi:10.1016/j.cogbrainres.2004.02.001.

7. Tobin, W. F., Wilson, R. I. & Lee, W. C. A. Wiring variations that enable and constrain neural computation in a sensory microcircuit. Elife (2017) doi:10.7554/eLife.24838.

8. Hiesinger, P. R. & Hassan, B. A. The Evolution of Variability and Robustness in Neural Development. Trends in Neurosciences (2018) doi:10.1016/j.tins.2018.05.007.

9. Waddington, C. H. Canalization of development and the inheritance of acquired characters. Nature (1942) doi:10.1038/150563a0.

10. Debat, V. Symmetry is beauty - Or is it? the rise and fall of fluctuating asymmetry. Medecine/Sciences (2016) doi:10.1051/medsci/20163208028.

11. Ayroles, J. F. et al. Behavioral idiosyncrasy reveals genetic control of phenotypic variability. Proc. Natl. Acad. Sci. U. S. A. (2015) doi:10.1073/pnas.1503830112.

12. Honegger, K. & de Bivort, B. Stochasticity, individuality and behavior. Current Biology (2018) doi:10.1016/j.cub.2017.11.058.

13. Pascual, A., Huang, K.-L., Neveu, J. & Préat, T. Brain asymmetry and long-term memory. Nature (2004) doi:10.1038/427605a.

14. Andriatsilavo, M. & Hassan, B. When brain development shapes individual behavior. Medecine/Sciences (2020) doi:10.1051/medsci/2020144.

15. Rihani, K. & Sachse, S. Shedding Light on Inter-Individual Variability of Olfactory Circuits in Drosophila. Front. Behav. Neurosci. 16, (2022).

16. Hassan, B. A. et al. Atonal regulates neurite arborization but does not act as a proneural gene in the Drosophila brain. Neuron (2000) doi:10.1016/S0896-6273(00)81059-4.

17. Zheng, X. et al. Baboon/dSmad2 TGF-β signaling is required during late larval stage for development of adult-specific neurons. EMBO J. (2006) doi:10.1038/sj.emboj.7600962.

18. Srahna, M. et al. A signaling network for patterning of neuronal connectivity in the Drosophila brain. PLoS Biol. (2006) doi:10.1371/journal.pbio.0040348.

19. Zschätzsch, M. et al. Regulation of branching dynamics by axon-intrinsic asymmetries in Tyrosine Kinase Receptor signaling. Elife (2014) doi:10.7554/elife.01699.

20. Agi, E., Kulkarni, A. & Hiesinger, P. R. Neuronal strategies for meeting the right partner during brain wiring. Current Opinion in Neurobiology (2020) doi:10.1016/j.conb.2020.01.002.

21. Han, C., Jan, L. Y. & Jan, Y. N. Enhancer-driven membrane markers for analysis of nonautonomous mechanisms reveal neuron-glia interactions in Drosophila. Proc. Natl. Acad. Sci. U. S. A. (2011) doi:10.1073/pnas.1106386108.

22. Langen, M. et al. The Developmental Rules of Neural Superposition in Drosophila. Cell (2015) doi:10.1016/j.cell.2015.05.055.

23. Özel, M. N., Langen, M., Hassan, B. A. & Hiesinger, P. R. Filopodial dynamics and growth cone stabilization in Drosophila visual circuit development. Elife (2015) doi:10.7554/eLife.10721.

24. Chan, Y. B., Alekseyenko, O. V. & Kravitz, E. A. Optogenetic control of gene expression in drosophila. PLoS One (2015) doi:10.1371/journal.pone.0138181.

25. Riedl, J. et al. Lifeact: A versatile marker to visualize F-actin. Nat. Methods (2008) doi:10.1038/nmeth.1220.

26. Grieder, N. C., bde Cuevas, M. & Spradling, A. C. The fusome organizes the microtubule network during oocyte differentiation in Drosophila. Development (2000) doi:10.1242/dev.127.19.4253.

27. Dumoulin, A., Zuñiga, N. R. & Stoeckli, E. T. Axon guidance at the spinal cord midline—A live imaging perspective. J. Comp. Neurol. (2021) doi:10.1002/cne.25107.

28. Moreland, T. & Poulain, F. E. To Stick or Not to Stick: The Multiple Roles of Cell Adhesion Molecules in Neural Circuit Assembly. Front. Neurosci. 16, 1–16 (2022).

29. Li, T. et al. Cellular Bases of Olfactory Circuit Assembly Revealed by Systematic Time-Lapse Imaging. SSRN Electron. J. (2021) doi:10.2139/ssrn.3839760.

30. Bastmeyer, M. & O’Leary, D. D. M. Dynamics of target recognition by interstitial axon branching along developing cortical axons. J. Neurosci. (1996) doi:10.1523/jneurosci.16-04-01450.1996.

31. Bovolenta, P. & Mason, C. Growth cone morphology varies with position in the developing mouse visual pathway from retina to first targets. J. Neurosci. 7, (1987).

32. Meltzer, H. & Schuldiner, O. With a little help from my friends: how intercellular communication shapes neuronal remodeling. Current Opinion in Neurobiology (2020) doi:10.1016/j.conb.2020.01.018.

33. Jang, K. J. et al. Two distinct filopodia populations at the growth cone allow to sense nanotopographical extracellular matrix cues to guide neurite outgrowth. PLoS One 5, (2010).

34. Turney, S. G. et al. Variation and selection in axon navigation through microtubuledependent stepwise growth cone advance. bioRxiv (2020).

35. Hassan, B. A. & Hiesinger, P. R. Beyond Molecular Codes: Simple Rules to Wire Complex Brains. Cell (2015) doi:10.1016/j.cell.2015.09.031.

36. Hiesinger, P. R. The Self-Assembling Brain. The Self-Assembling Brain (2021). doi:10.2307/j.ctv191kwz2.

## Methods References

1. B. A. Hassan, N. A. Bermingham, Y. He, Y. Sun, Y. N. Jan, H. Y. Zoghbi, H. J. Bellen, Atonal regulates neurite arborization but does not act as a proneural gene in the Drosophila brain. Neuron (2000), doi:10.1016/S0896-6273(00)81059-4.

2. C. Han, L. Y. Jan, Y. N. Jan, Enhancer-driven membrane markers for analysis of nonautonomous mechanisms reveal neuron-glia interactions in Drosophila. Proc. Natl. Acad. Sci. U. S. A. (2011), doi:10.1073/pnas.1106386108.

3. A. Nern, B. D. Pfeiffer, G. M. Rubin, Optimized tools for multicolor stochastic labeling reveal diverse stereotyped cell arrangements in the fly visual system. Proc. Natl. Acad. Sci. U. S. A. 112, E2967–E2976 (2015).

4. G. Struhl, I. Greenwald, Presenilin is required for activity and nuclear access of notch in drosophila. Nature (1999), doi:10.1038/19091.

5. S. Weinberger, M. P. Topping, J. Yan, A. Claeys, N. De Geest, D. Ozbay, T. Hassan, X. He, J. T. Albert, B. A. Hassan, A. Ramaekers, Evolutionary changes in transcription factor coding sequence quantitatively alter sensory organ development and function. Elife. 6 (2017), doi:10.7554/eLife.26402.

6. Y. B. Chan, O. V. Alekseyenko, E. A. Kravitz, Optogenetic control of gene expression in drosophila. PLoS One (2015), doi:10.1371/journal.pone.0138181.

7. N. C. Grieder, M. de Cuevas, A. C. Spradling, The fusome organizes the microtubule network during oocyte differentiation in Drosophila. Development (2000), doi:10.1242/dev.127.19.4253.

8. A. Krejčí, F. Bernard, B. E. Housden, S. Collins, S. J. Bray, Direct response to notch activation: Signaling crosstalk and incoherent logic. Sci. Signal. 2 (2009), doi:10.1126/scisignal.2000140.

9. M. N. Özel, M. Langen, B. A. Hassan, P. R. Hiesinger, Filopodial dynamics and growth cone stabilization in Drosophila visual circuit development. Elife (2015), doi:10.7554/eLife.10721.

10. K. T. Ngo, I. Andrade, V. Hartenstein, Spatio-temporal pattern of neuronal differentiation in the Drosophila visual system: A user’s guide to the dynamic morphology of the developing optic lobe. Dev. Biol. (2017), doi:10.1016/j.ydbio.2017.05.008.

11. M. Langen, E. Agi, D. J. Altschuler, L. F. Wu, S. J. Altschuler, P. R. Hiesinger, The Developmental Rules of Neural Superposition in Drosophila. Cell (2015), doi:10.1016/j.cell.2015.05.055.

12. K. White, D. R. Kankel, Patterns of cell division and cell movement in the formation of the imaginal nervous system in Drosophila melanogaster. Dev. Biol. 65 (1978), doi:10.1016/0012-1606(78)90029-5.

13. A. Dumoulin, N. R. Zuñiga, E. T. Stoeckli, Axon guidance at the spinal cord midline— A live imaging perspective. J. Comp. Neurol. (2021), doi:10.1002/cne.25107.

14. Sagar F. Pröls, C. Wiegreffe, M. Scaal, Communication between distant epithelial cells by filopodia-like protrusions during embryonic development. Dev. (2015), doi:10.1242/dev.115964.

15. N. H. Wilson, E. T. Stoeckli, In ovo electroporation of miRNA-based plasmids in the developing neural tube and assessment of phenotypes by DiI injection in open-book preparations. J. Vis. Exp. (2012), doi:10.3791/4384.

16. V. Hamburger, H. L. Hamilton, A series of normal stages in the development of the chick embryo. J. Morphol. (1951), doi:10.1002/jmor.1050880104.

17. J. Schindelin, I. Arganda-Carreras, E. Frise, V. Kaynig, M. Longair, T. Pietzsch, S. Preibisch, C. Rueden, S. Saalfeld, B. Schmid, J. Y. Tinevez, D. J. White, V. Hartenstein, K. Eliceiri, P. Tomancak, A. Cardona, Fiji: An open-source platform for biological-image analysis. Nat. Methods (2012),, doi:10.1038/nmeth.2019.

